# The predictive power of the microbiome exceeds that of genome-wide association studies in the discrimination of complex human disease

**DOI:** 10.1101/2019.12.31.891978

**Authors:** Braden T Tierney, Yixuan He, George M Church, Eran Segal, Aleksandar D Kostic, Chirag J Patel

## Abstract

Over the past decade, studies of the human genome and microbiome have deepened our understanding of the connections between human genes, environments, microbes, and disease. For example, the sheer number of indicators of the microbiome and human genetic common variants associated with disease has been immense, but clinical utility has been elusive. Here, we compared the predictive capabilities of the human microbiome versus human genomic common variants across 13 common diseases. We concluded that microbiomic indicators outperform human genetics in predicting host phenotype (overall Microbiome-Association-Study [MAS] area under the curve [AUC] = 0.79 [SE = 0.03], overall Genome-Wide-Association-Study [GWAS] AUC = 0.67 [SE = 0.02]). Our results, while preliminary and focused on a subset of the totality of disease, demonstrate the relative predictive ability of the microbiome, indicating that it may outperform human genetics in discriminating human disease cases and controls. They additionally motivate the need for population-level microbiome sequencing resources, akin to the UK Biobank, to further improve and reproduce metagenomic models of disease.

## Main Text

We have come to know the human microbiome, or metagenome, as our “second genome,”(Grice and Segre 2012) and, indeed, the human genome and metagenome share many features (Figure 1A). Both are, at their cores, networks of genes that affect host health and vary across human populations (Thingholm et al. 2019; Morgan et al. 2012; Vatanen et al. 2016; Minot and Willis 2019; Boyle, Li, and Pritchard 2017). However, there are striking dissimilarities as well, notably in terms of scale (∼20,000 human genes vs. tens of millions of microbial genes across thousands of species), the transient nature of human microbiota, and the sensitivity of the microbiome to the host’s environment(Rothschild et al. 2018; Tierney et al. 2019; Moraes and Góes 2016; Theriot et al. 2014) We hypothesized that since the microbiome may be viewed as a hub that captures host environmental factors, microbial features may provide sources of variation not captured by static host common variants. Therefore, a natural question arises when reflecting upon our “first” and “second” genomes: how are microbiome and human genome different (or complementary) in their ability to characterize host phenotype, and which one is superior at doing so overall?

**Figure 1:**
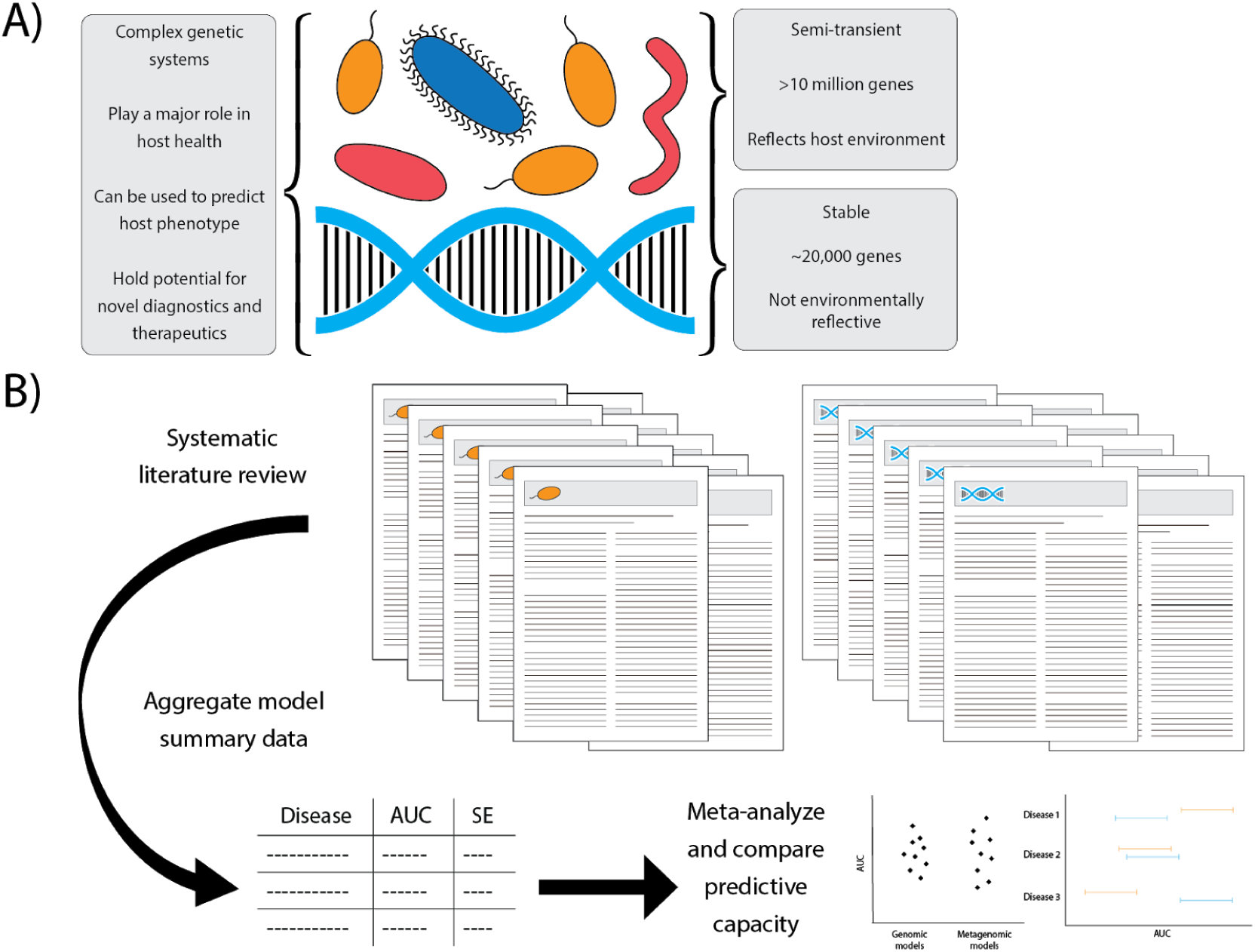
Overview. A) Key differences and similarities between the human genome and metagenome. B) Meta-analytic process for comparing predictive capacities of human microbiome vs. human genome.

Answering these questions would have vast implications for our understanding of the genome’s and microbiome’s respective diagnostic and therapeutic potentials. Rothschild and colleagues have documented a way to measure the total variance explained of metagenomic indicators in complex traits called the “microbiome-association-index” (*b*^*2*^) (Rothschild et al. 2018). This measure quantifies the total variance explained between microbiota and host phenotype after accounting for genetic variation and is analogous to heritability (*h*^*2*^) (Peter M Visscher, Hill, and Naomi R Wray 2008), which describes variation in a phenotype attributable to genetics.

However, the comparison between the predictive capability of metagenomics and common variants of the human genome has been only anecdotal to date. Here, to begin to address this gap, we aimed to quantify differences in the ability of the metagenome versus common variants of the human genome to predict the presence of specific host phenotypes in case-control studies.

We meta-analyzed the predictive capacity of the metagenome from disparate investigations (Figure 1B).(Kavvoura and Ioannidis 2008) A number of teams over the past decade have associated host phenotype from either metagenomic or human single nucleotide polymorphism (SNP) data via Microbial-Association-Studies (MAS) or Genome-Wide Association Studies (GWAS) respectively.(Timpson et al. 2018) For the purposes of this investigation, an MAS is a study that associates microbiome features, like species abundance, with disease presence in a case-control study. A GWAS aims to associate millions of common genetic variants with disease task (for a full table of definitions relevant to this work, see Supp. Table 1). Common variants have a higher prevalence in the population (1-10%). Current GWAS arrays measure on the order of millions of common variants; however, the thousands of species and metabolic MAS elements used in our aggregated predictors are not constrained by prevalence.

**Table 1:**
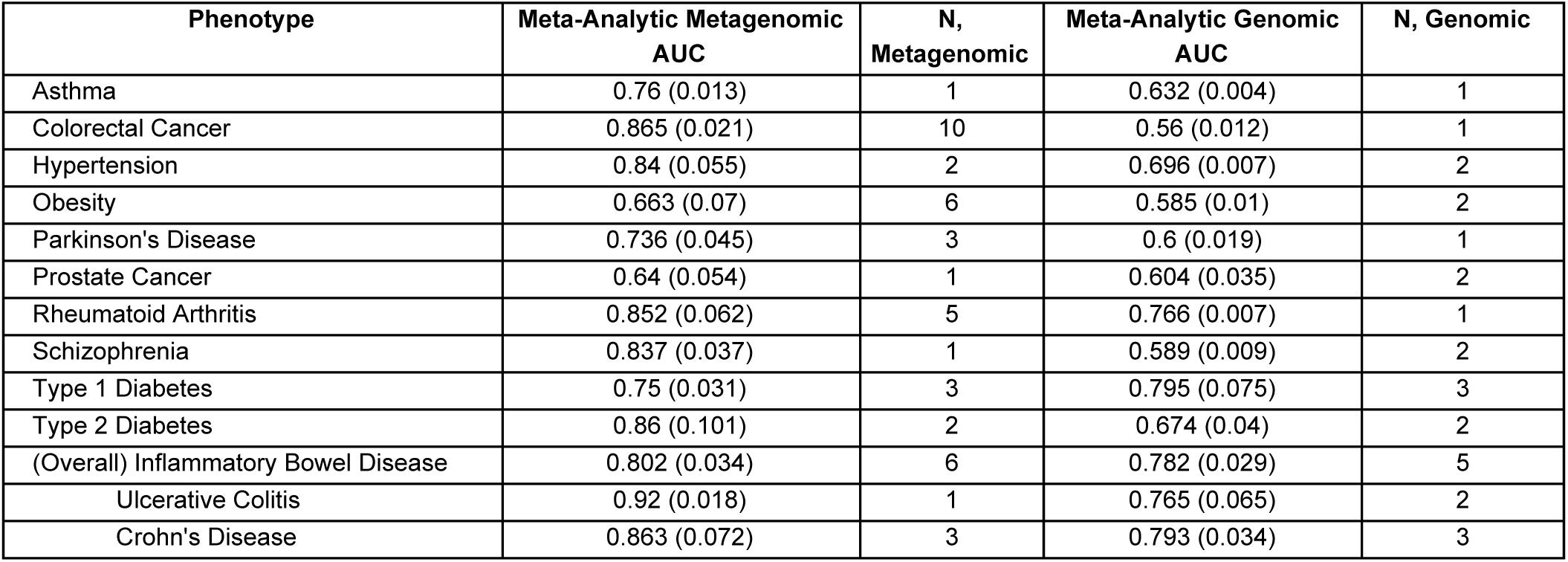
Meta-analysis of genomic vs. metagenomic predictors. Standard errors are in parentheses, followed by the number of models used for a given disease.

While MAS and GWAS are important to find individual variants or microbial genes associated with disease, the aggregate total variants they identify may be clinically informative in their predictive utility, for example in exceptional individuals identified in a “polygenic risk score” (Khera et al. 2018). Different groups have published on the same diseases in independent cohorts and populations, assessing microbial/SNP predictive capability with area under the receiver operator curve (AUC). An AUC of 1.0 implies perfect classification of disease cases and controls, whereas an AUC of 0.5 means the model is performing no better than random choice. While individual studies may be biased by small sample sizes or methodological variation, meta-analyzing across independent studies allow investigators to synthesize evidence.(DerSimonian and Laird 2015)

Naturally, GWAS is bound to outperform the microbiome for certain phenotypes (e.g., sex, height) and vice versa (e.g., *Clostridium difficile* infection). Therefore, we chose to focus on exclusively complex diseases known to arise from a mixture of environmental and genetic factors, systematically searching the literature for microbiome and SNP-based genomic studies that built predictive models host disease. In total, we identified 30 publications, 47 distinct MASs and 24 GWASs (Supp. Table 2).

We found that metagenomic predictors outperform human common genomic variants in classifying hosts based on phenotype (Table 1, Figure 2). The standard-error weighted average AUC for the aggregated metagenomic predictors was 0.79 (SE = 0.03), compared to 0.67 (SE = 0.02) for GWAS (Figure 2). The phenotype with the broadest discrepancy between meta-analytic metagenomic and genomic predictors was colorectal cancer (metagenomic AUC = 0.83 [SE = 0.021], genomic AUC = 0.56 [SE = 0.012]). The only disease where GWAS outperformed the metagenome in our meta-analytic estimates was Type 1 Diabetes (metagenomic AUC = 0.75 [SE = 0.031], genomic AUC = 0.80 [SE = 0.075]).

**Figure 2:**
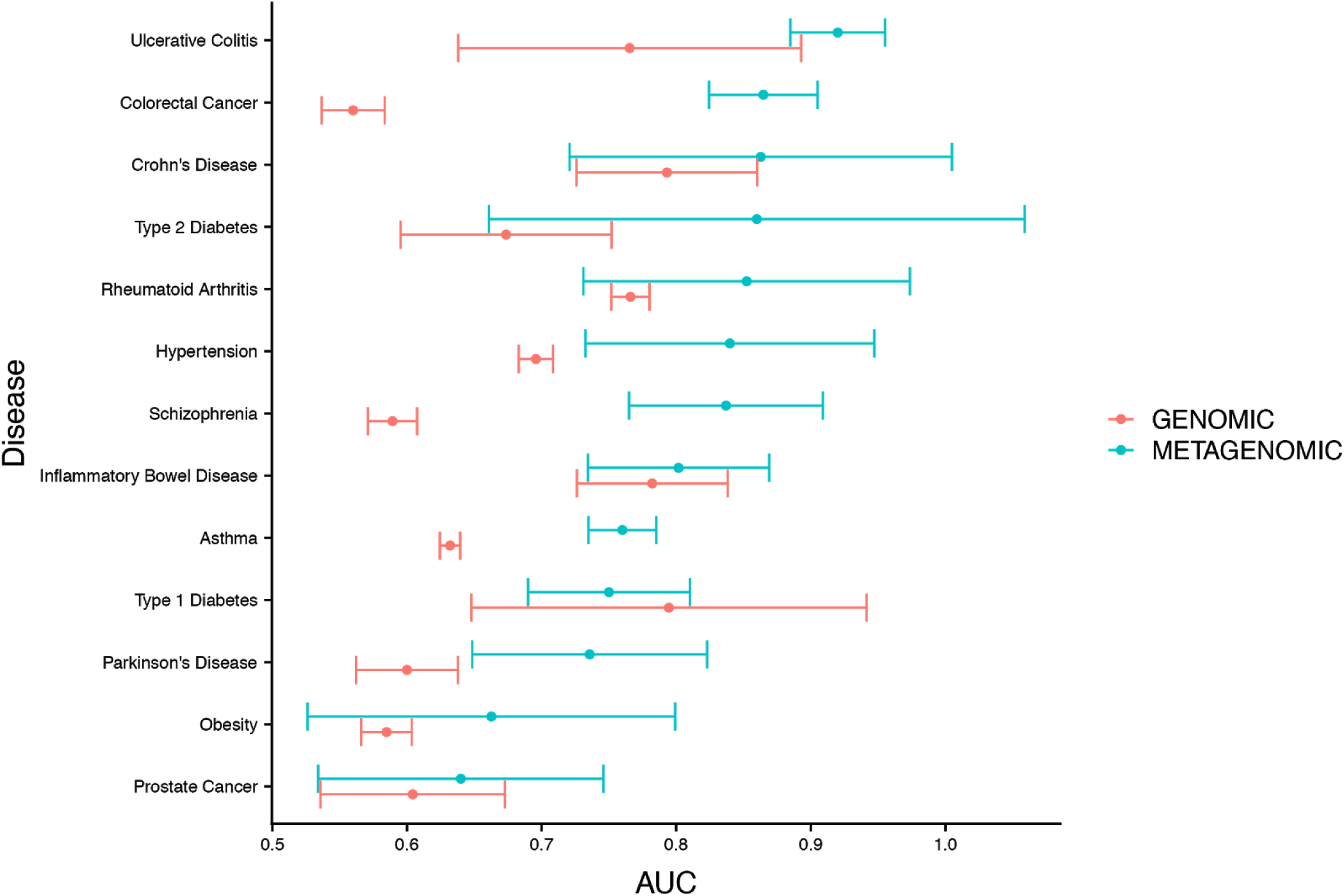
Results from literature-review-based meta-analysis of genomic vs. metagenomic predictors.

Findings from meta-analyses may be biased due to selective publication biases. We found that both metagenomic and genomic studies exhibit publication bias for both the metagenomic and genomic studies (metagenomic p-value of p = 0.0003, genomic p-value of p < 0.0001), with the latter demonstrating increased slightly funnel plot asymmetry (Supp Fig 2).

We used alternative measures to compare MAS and GWAS to complement our systematic review. First, we compared, using published estimates,(Rothschild et al. 2018; “Heritability of >4,000 Traits & Disorders in UK Biobank” n.d.) the microbiome-association-index (*b*^*2*^) to estimates of common variant heritability (*h*^*2*^) for a series of quantitative traits (Supp Table 3). We found *b*^*2*^ to explain more variance in phenotype than *h*^*2*^ for 6/8 traits, with the only exceptions being height and total cholesterol. As a second method of comparison, we analyzed cohort data from the UK Biobank(Sudlow et al. 2015) to build our own predictors of disease using GWAS-significant common variants (Supp Table 3). We found the literature-based metagenomic estimators to universally outperform the genomic AUCs (overall genomic AUC = 0.57, SE = 0.0091). Finally, we estimated the maximum AUC (Max_AUC_G) a set of SNPs could theoretically generate for a given disease prevalence and heritability.(Wray et al. 2010) The theoretical maximum AUCs assume that the association sizes for all variants have been identified. Given that each individual variant explains a small amount of disease risk, large sample sizes are required to extract signal from noise. We found that the microbiome literature-based estimates approached the theoretical maximum genetic AUCs. (Supp Table 3).

In short, based on an exhaustive search through a decade of literature and a range of measures (AUC, h^2^, and b^2^), we conclude that the metagenome is a better predictor of human disease status than common variants of the human genome. We additionally have identified that published metagenomic predictors are approaching the theoretical limit of predictive power for genomic common variants – a feat not even attained by any of the genomic predictors in our literature review and potentially impossible to attain with the largest sample sizes. The major limitations of our study arise from 1) the available literature and its associated publication bias, 2) the predominant reliance of metagenomic vs. genomic studies on different tools for prediction with assumptions on the mode of association (linear regression and non-linear random forests, respectively), 3) lack of consideration of the changing nature of the microbiome, 4) the more established methodology of GWAS, specifically regarding the use of validation cohorts in most studies (this is not universally the case for the microbiome), and relatedly, 5) the comparably lower sample size in microbiome studies may have led to “overfitting” in the initial studies(Topçuoğlu et al. 2019) and susceptibility to “Winner’s Curse” bias, phenomena that are exemplified by inflated initial and non-replicated findings (Ioannidis, Thomas, and Daly 2009). Furthermore, all of these predictors (GWAS and MAS) may not be as useful as existing tools for clinical risk prediction, such as the Framingham Risk Score (Wang et al. 2012). If MAS predictors are deployed en masse, what can be done after reporting to patients their microbial disease prediction? Currently, not much. The path to medical decision consists of many more steps beyond evaluating AUCs, such as understanding if they are informative of intervention or even if they patients fare better after them. New approaches will be needed to assess MAS predictors in the context of existing predictors and decision pathways (Bossuyt et al. 2006).

A way to address these problems holistically is to invest in population-scale and longitudinal resources for the microbiome, as we have for human genetics (i.e. the UK Biobank). Increasing patient observation window lengths and sample sizes from thousands (the scale of the Human Microbiome Project(Human Microbiome Project Consortium 2012)) to hundreds of thousands (the scale of the UK Biobank) will enable researchers to test the predictive limits of the human microbiome. The difference in predictive performance between MAS and GWAS is further amplified when considering this difference in resource allocation – with increased investment in large-scale microbiome resources, we hypothesize the gap between MAS and GWAS AUCs will further increase.

It is worth investigating further the optimal data types for predicting host phenotype from metagenomic/genomic data. For example, the microbiome can be viewed in terms of microbial species, metabolic pathways, individual genes, or even in individual SNPs. The metagenomic space contains millions of genes(Tierney et al. 2019) and encompasses complex structural variation on top of simple single nucleotide polymorphisms (SNPs). Host genetic features, on the other hand, are not limited to exclusively the common variants measured on a GWAS chip, which only measures the ∼0.1% of where the human genome contains variation(1000 Genomes Project Consortium et al. 2015). Feasibly, GWAS AUCs could also increase if they incorporated rare features or other genomic elements (i.e. insertions-deletions or copy-number variants). While we have shown here that on average metagenomics outperforms SNP-based arrays, future work should include doing comparisons between all of these different data types, as they will likely perform differently in terms of their predictive capability.

Finally, it is worth noting that predictive performance does not imply causality. Currently, determining causality in the human microbiome is challenging and addressed largely through model system experimentation. Identifying causal variants is less intense as the human genetic variants are static and randomly assorted(Ioannidis et al. 2009). Further, in genetic studies, Issues of confounding via population stratification(Price et al. 2006) have largely been solved, but identifying confounders in dynamic variables are elusive(Manrai, Ioannidis, and Patel 2019). Therefore, while the metagenome may outperform GWAS for predicting disease, determining causal relationships between the microbiome and disease is potentially more difficult.

Our work is only a gateway study; as we have noted, there is a need for future efforts and methodological development. With increased metagenomic sequencing, we will be able to further this work and test the limits of the microbiome in its ability to characterize all manners of host phenotypes. Hopefully, a day is coming soon where metagenomic resources are merged with genomics and clinical histories on par, in terms of scale, with those that currently exist for genomics. Only then will we seriously be able to move the microbiome into the clinic *en masse*.

---

**Supplementary Figure 1:**
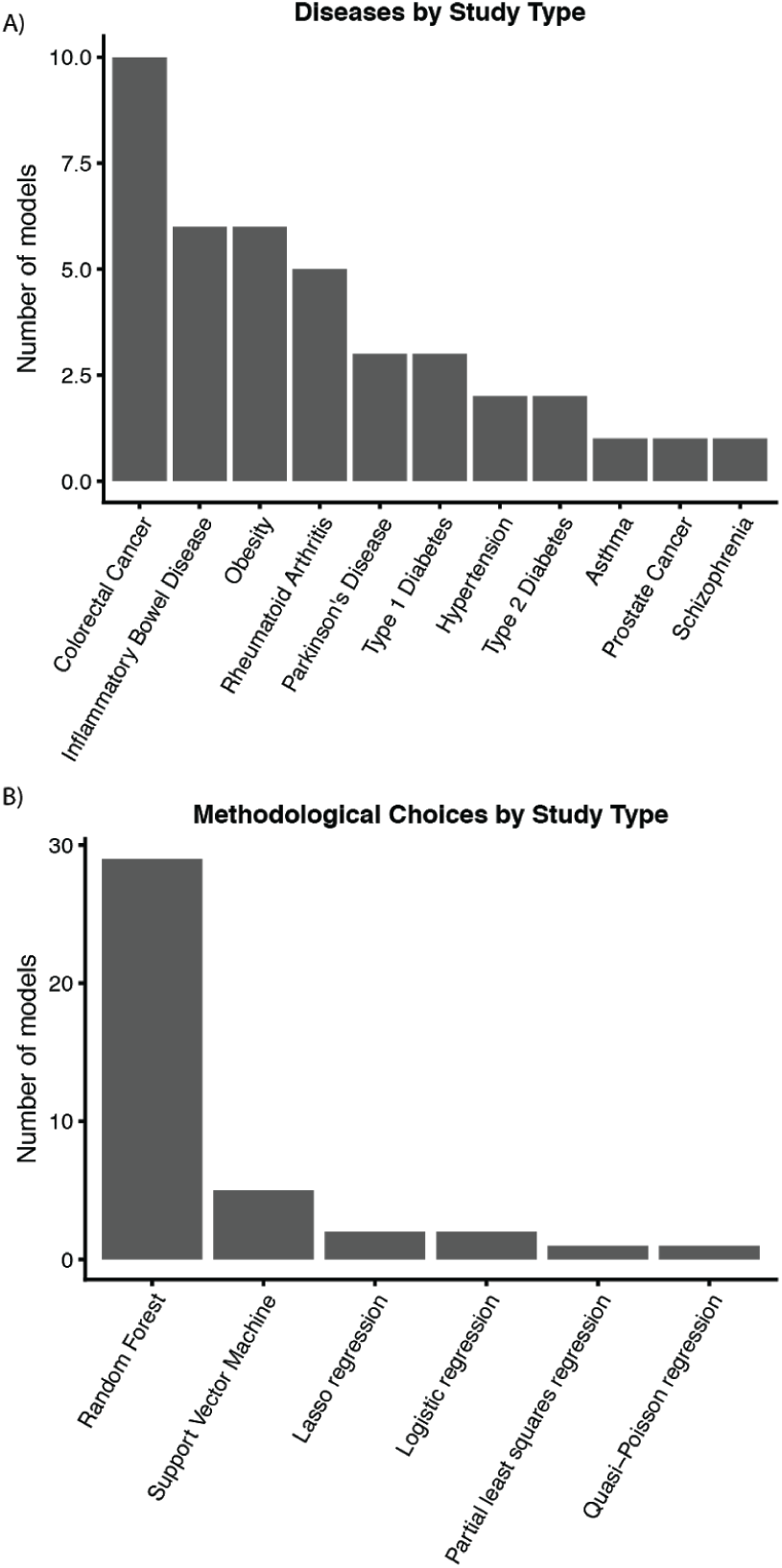
Summarized literature review in terms of A) number of models per disease aggregated and B) distribution of prediction methods.

**Supplementary Figure 2:**
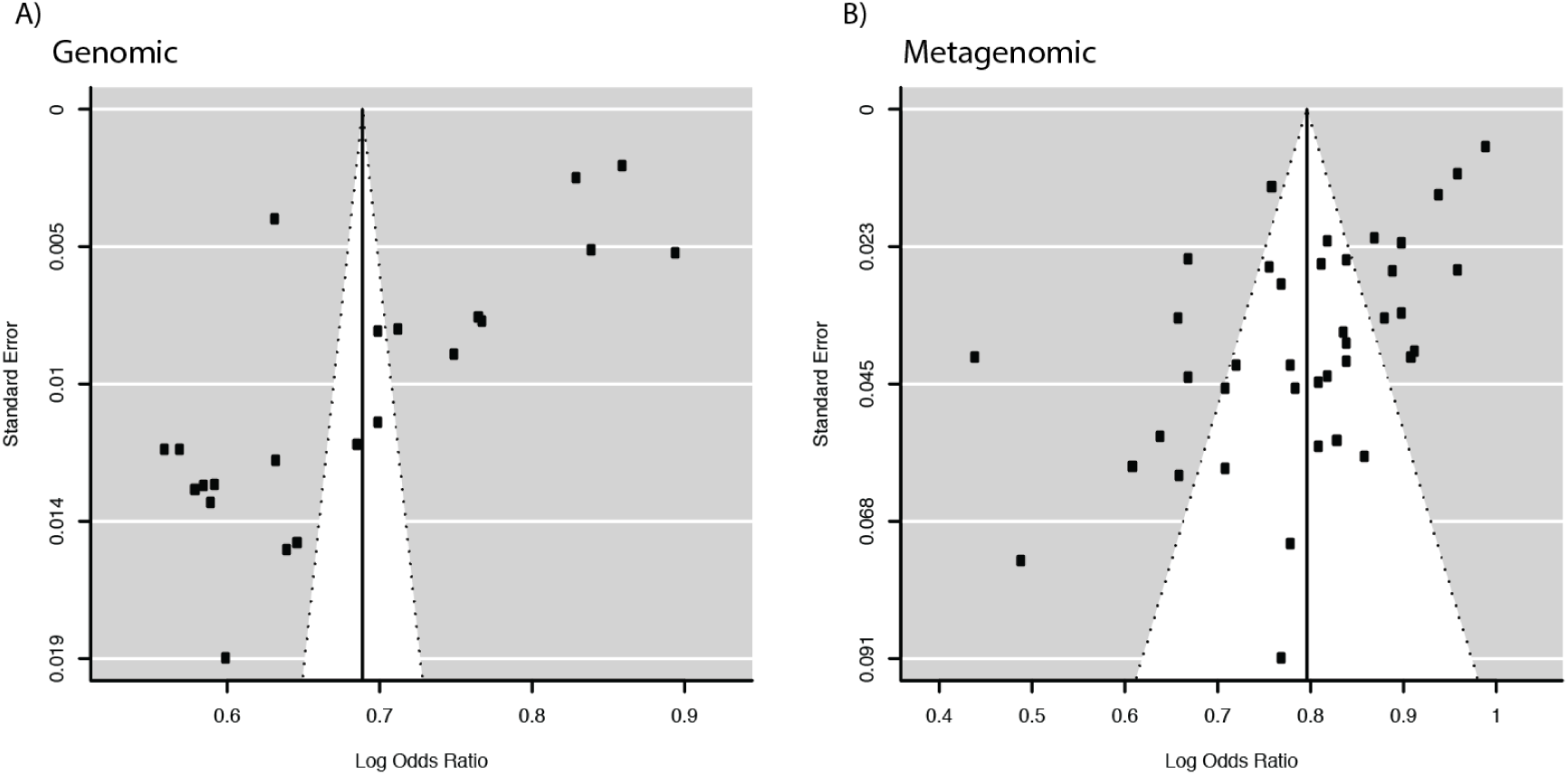
Funnel plots of AUCs from literature review for A) genomic predictors and B) metagenomic predictors.

Supplementary Table 1: Table of definitions used in study. Supplementary Table 2: Literature review process and raw findings.

Supplementary Table 3: Alternative estimations of common variant AUCs, including AUC_PRS estimation, *h*^*2*^ versus *b*^*2*^ comparison, and Max_AUC_G estimation.

Supplementary Table 4: Relevant information for phenotyping from the UK Biobank and mapping ICD9/10 to PHEWAS codes.

## Methods

### Overview of the approach

We aimed to compare the predictive capacity of the microbiome and the human genome via a meta-analytic approach. We identified studies that predicted disease state (versus non-disease controls), with the input being either common human genetic variants or metagenomic data. By performing a systematic meta-analysis, we aimed to minimize variation due to individual study bias or batch effects (i.e. due to model choice, cohort characteristics, or other methodological variation).

We selected studies that 1) clearly reported their methods and sample sizes and 2) used area under the receiver-operator (sensitivity/specificity) curve (AUC) as their summary statistics measure for classification accuracy. AUC is used to quantify model performance regarding discriminating between the two groups. An AUC of 1 means a model is perfectly separating cases and controls, whereas 0.5 is no better than randomly guessing.

As a further point of comparison, we additionally utilized three alternative methods for computing the maximum possible AUC genomic information could provide in predicting disease incidence. Extensive research in the genomic space has identified a link between disease heritability – the amount of variation in disease explainable by genetic factors – and AUC. Therefore, using prior published methods, we were able to compute theoretical maximum AUC values using heritability estimates derived from GWAS common variants (referred to as Max_AUC_G). Second, we compared the “microbiome-explainability” statistic (b^2^)(Rothschild et al. 2018) to heritability. Third, we built our own predictors for our traits of interest using the UK Biobank and reported the output AUCs.

### Metagenomic study aggregation

We performed a systematic literature review with the aim of identifying any study that sought to associate host phenotype with microbiome contents. We searched broadly on PubMed for studies associated with the terms “(microbiome wide association and human) or (metagenome wide association study and human),” identifying 507 publications. We read the abstract and titles of each of these to identify if they were likely to report an AUC or perform any manner of prediction. We additionally performed a suite of narrower, disease-specific, searches on Google Scholar, trying to identify studies published on specific diseases. Our full search terms are described in Supplementary Table 1. We only selected studies that did not adjust for age, sex, or any demographic variables, instead only using microbiome feature to predict phenotype.

### Genomic study aggregation

After identifying the set of diseases that had metagenomic-predictors built for them, we sought to identify a set of publications that build predictors the same phenotypes. We used targeted searches for each disease in Google Scholar (i.e. “genomic predictor AUC asthma”) that are reported in Supplementary Table 1. We additionally carried out a broad, systematic search in PubMed across 76 studies using the search terms (“complex trait disease prediction genomic snp “). We only selected studies that did not adjust for age, sex, or any demographic variables, instead only using common variants to predict phenotype.

### Meta-analysis of published AUCs

We extracted AUCs and sample sizes for each metagenomic and genomic study and computed standard errors on them via the auctestr (https://www.rdocumentation.org/packages/auctestr) package in R. When possible, we only took AUCs generated from testing or validation cohorts. For our random-effects meta-analysis, we used the metafor(Viechtbauer 2010) package to compute standard-error-weighted averages of our AUCs within each disease.

### Phenotype ascertainment from the UK Biobank

We classified phenotypes based on self-reported data during an interview with a trained nurse as well as International Classification of Diseases (ICD-9 and ICD-10) diagnostic codes in the UK Biobank. We then used PHEWAS codes from the Phecode Map 1.2(Wu et al. 2018) to map ICD codes to phenotypes. The full list of PHEWAS to ICD code mappings are available in Supplementary Table 4. For a given PHEWAS code (i.e. Rheumatoid Arthritis), we identified all occurrences of its component ICD10 and ICD9 codes in the UK Biobank. We encoded individuals a 0 or 1, where 1 referred to the presence of disease. We counted the presence of any 1 relevant ICD9/10 code at any point for a given patient as the presence of that disease.

### Computing max genomic AUC (Max_AUC_G)

To compute the theoretical maximum genomic AUC values that could be achieved from common GWAS variants, we implemented a method (Wray et al. 2010) (https://github.com/kn3in/genRoc) that converts between heritability on the liability scale and AUC. We estimated disease heritability with BOLT-REML V2.3.4 (Loh et al. 2018). In order to capture predictive capability only deriving from common SNPs, we did not adjust for age or sex in running BOLT-REML. Afterwards we converted observed heritability to the liability scale using disease prevalence within the sample population and the global population. We estimated these with prevalances from the UK Biobank and World Health Organization, respectively.

### Computing genomic AUC (AUC_PRS)

We computed the genomic AUC from a polygenic risk score (PRS) built from common GWAS variants. To do this, we separated the UK biobank White European population (N=418,114) into three cohorts in a 1:1:2 ratio. We conducted a simple GWAS without any covariates in group A. Next, we calculate weights for each of the common variants using PRS-CS(Ge et al. 2019), which takes the GWAS summary statistics from group A and an external linkage disequilibrium (LD) reference panel to estimate effect sizes in group B. Using these weights, we calculated PRS’s for individuals in group C using PLINK(Chang et al. 2015) with its built in allelic scoring procedure (--score). We report the average of 100 AUC’s (bootstrapped 1000 iterations) calculated in sampled populations of balanced cases and controls. AUCs were calculated using the ‘auc’ function in the pROC(Robin et al. 2011) package.

